# Hierarchical modeling of the effect of pre-exposure prophylaxis on HIV in the US

**DOI:** 10.1101/285940

**Authors:** Renee Dale, Ying Chen, Hongyu He

**Affiliations:** Department of Experimental Statistics, Louisiana State University, Baton Rouge, Louisiana, United States of America; Department of Biological Sciences, Louisiana State University, Baton Rouge, Louisiana, United States of America; Fred Hutchinson Cancer Center, Washington; Department of Mathematics, Louisiana State University, Baton Rouge, Louisiana, United States of America

**Keywords:** HIV, Mathematical modeling, Differential equations, Pre-exposure prophylaxis, Rural healthcare, Behavior risk, Epidemiology

## Abstract

In this paper we present a differential equation model stratified by behavioral risk and sexual activity. Some susceptible individuals have higher rates of risky behavior that increase their chance of contracting the disease. Infected individuals can be considered to be generally sexually active or inactive. The sexually active infected population is at higher risk of transmitting the disease to a susceptible individual. We further divide the sexually active population into diagnosed or undiagnosed infected individuals. We define model parameters for both the national and the urban case. These parameter sets are used to study the predicted population dynamics over the next 5 years. Our results indicate that the undiagnosed high risk infected group is the largest contributor to the epidemic. Finally, we apply a preventative medication protocol to the susceptible population and observe the effective reduction in the infected population. The simulations suggest that preventative medication effectiveness extends outside of the group that is taking the drug (herd immunity). Our models suggest that a strategy targeting the high risk undiagnosed infected group would have the largest impact in the next 5 years. We also find that such a protocol has similar effects for the national as the urban case, despite the smaller sexual network found in rural areas.

## 2. Introduction

HIV is disease found across the world that causes autoimmune deficiency syndrome (AIDS). Drugs that treat HIV and prevent AIDS, called anti-retroviral therapies (ART), were first developed in 1985 [4]. However, the HIV infected population shows a steady growth rate [3, 5, 9]. PreP (pre-exposure prophylaxis) is a possible solution to the problem of the continuous increase in the population size. It was found that the rate of transmission of HIV was reduced by 44% when HIV-negative individuals took ART medications as chemoprophylaxis [14]. It is currently estimated that approximately 135,000 individuals in the US are prescribed chemoprophylaxis [12]. To understand what an effective prevalence would be, we first need to understand the composition of the susceptible population. Previous research has found that the chance of transmitting HIV is largely dependent on partner number [1, 2]. It is well known that due to higher population density, the urban and rural populations do not have the same dynamics. We use Detroit to look at the urban population dynamics. This was inspired by the recent paper which estimated the transmission rates [8]. The Detroit diagnosis and mortality rate is much higher than the national case [9, 13].

In this paper we establish two susceptible populations corresponding to an average-density sexual network (national) and high-density sexual network (urban). We then stratify the susceptible population by high and low risk sexually active individuals. Previously it was found that the effect of susceptible risk cannot be ignored when considering infected dynamics in the US [11]. We consider the susceptible population to be about 10 times larger than the infected population, a conservative estimate since the primary groups affected by the HIV epidemic are MSM and IV drug users [9]. Previous research estimates the size of the high risk population to be about 10%, and we use this number to get a conservative estimate of the high risk susceptible population [2]. By considering these two susceptible populations, we seek to shed light on the method to optimally prevent further increase in the magnitude of the epidemic.

Risky behavior affects transmission events in two ways. High risk susceptibles are more likely to engage in high-risk sexual activities, and they are more likely to do this with a higher partner number. On the infected side, there is the chance that high risk individuals may not be aware of their status. This correlates with an increased likelihood of risky behavior, since risky behavior within this population is reduced by approximately 50% after diagnosis [1] When considering the effectiveness of PreP for a given population of susceptibles, we must also consider the many factors involved in those susceptibles contracting HIV. We model the infected population by stratification along awareness of their HIV status and their risk level. We further consider that, regardless of awareness level, some infected individuals may be largely sexually inactive. The transmission of HIV for this group of individuals we consider to be due to chance or rare sexual encounters. Therefore we consider the risk per encounter of HIV transmission is similar between low risk diagnosed and undiagnosed individuals. In our model, the infected population is stratified into three levels: high risk diagnosed, high risk undiagnosed, and low risk infected.

We use our model to look at the effect of a targeted protocol on the rate at which susceptible individuals contract HIV. Our simulations allow us to calculate the optimal target population for PreP usage based on the projected 5 year population dynamics in high and low risk susceptibles for urban or national (rural and urban) conditions. Our model suggests that targeting the high risk susceptible population is always more effective. Although this result is intuitive, we also find that such a protocol is equally effective at the national and urban levels. This demonstrates the unintuitive effect of targeting small sexual networks in the prevention of HIV transmission in the United States.

## 3. Model

To construct our model (Fig. 2), we first separate the susceptible population into high risk *S*_*h*_ and low risk *S*_*l*_. We use *P* (*t*) to describe the growth rate of the susceptibles, estimated to be roughly the population growth rate in general. We consider new susceptibles to be relatively low risk, and movement into the high risk category is modeled using *g*(*t*).

**Figure 1.**
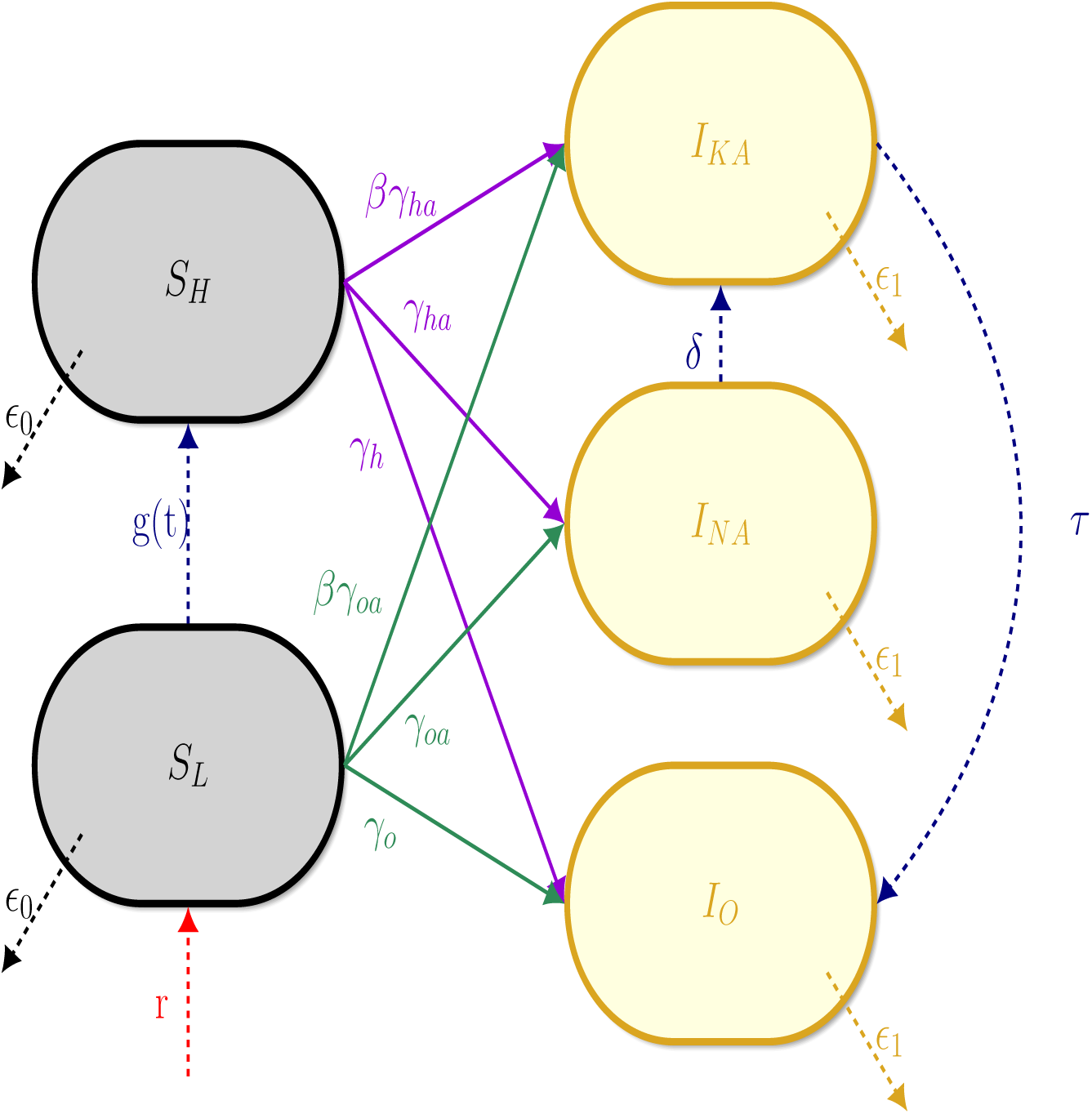
Conceptual model of the system of susceptibles *S*, infected *I*, with movement described by the parameters from Table 1.

**Figure 2.**
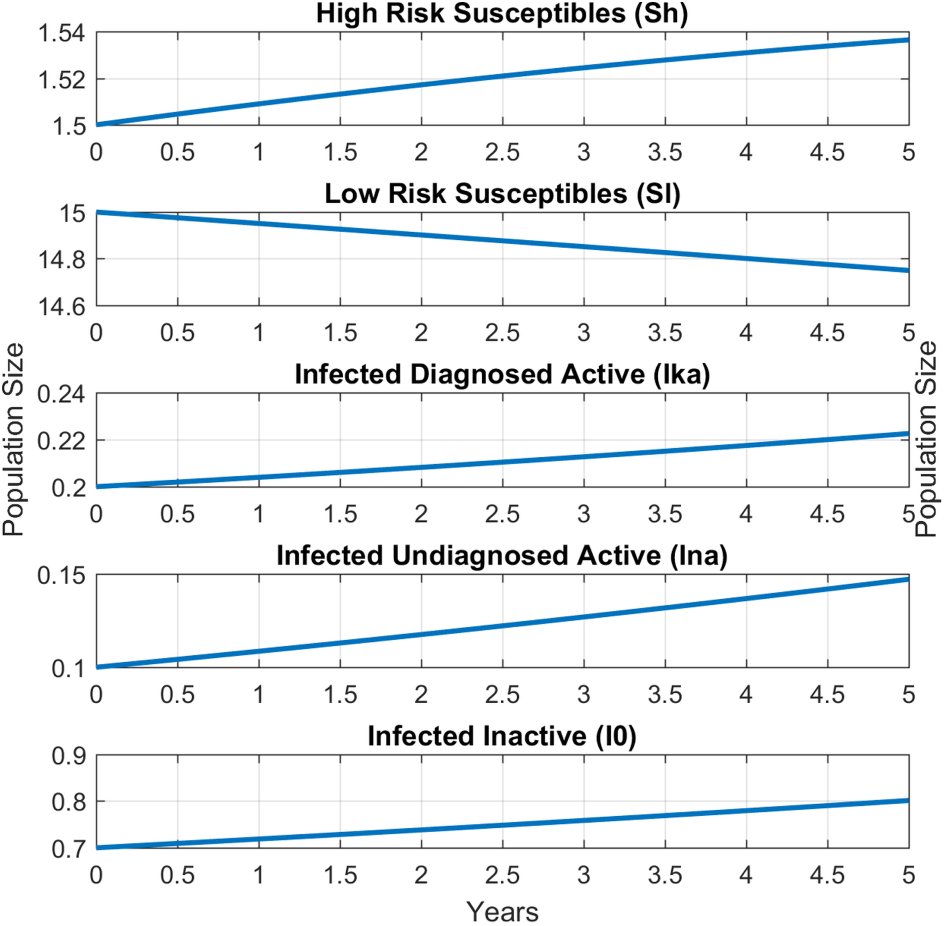
Model simulation of national HIV population dynamics. Parameter estimates in Table 1

The infected population is stratified by both sexual activity and diagnosis status. High risk undiagnosed individuals are designated as *I*_*na*_ (infected not aware active), and diagnosed individuals as *I*_*ka*_ (infected knowing active). Movement from undiagnosed to diagnosed is modeled using the diagnosis rate *δ*. Data suggests that most newly infected individuals are diagnosed within one year of contracting the disease [3]. We use *θ* ∈ [0,1] to distribute these individuals between diagnosed and undiagnosed populations proportionally. We consider low risk individuals’ chance of transmitting the disease to be indistinguishable by diagnosis status. Both undiagnosed and diagnosed low risk populations are pooled into a single population *I*_*o*_.

Transmission rates are modified according to the relative risk of each group. Infected individuals who are aware are considered to have significant reduction in transmissibility due to reduction in risky behavior, ART usage, and other, psychological factors [1, 2]. Non-active individuals have a very low chance of transmitting the disease, reflective of their low partner number. Death rates of all groups are estimated using data [9, 5]. The following figure illustrates the population dynamics in our model.

We obtain the following system of equations to describe these dynamics:

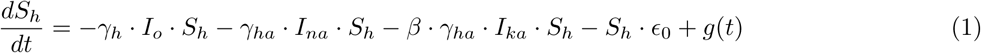

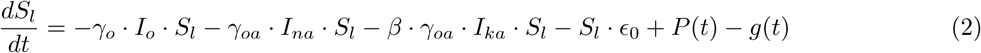

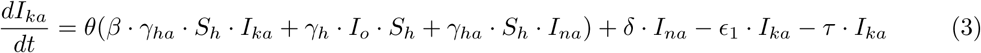

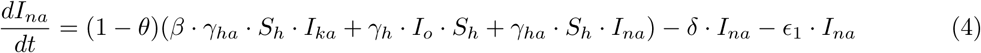

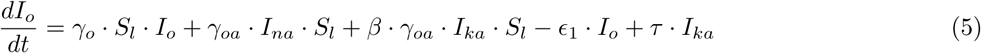

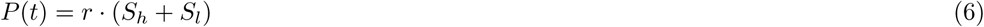

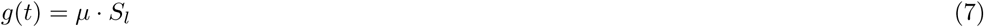

## 4. National population dynamics

Current estimates put the size of the high risk populations at 10% of the population [2]. The susceptible population can be estimated as 8 times larger than the total infected population. CD4 estimates of the undiagnosed population put it at about 20% of the total infected population [3].

The highest transmission rate is between high risk susceptibles and high risk infected who are undiagnosed (*γ*_*h*_). Previously it was found that high risk individuals are more likely to get diagnosed in the early stages of the disease, where transmission rate is the highest [10]. We estimate this using early stage transmission rates from [8]. The transmission rate for high risk susceptibles with high risk infecteds who are aware of their HIV status is reduced by 50% (*β*) [1]. Contact between high risk susceptibles and low risk infecteds is estimated using chronic HIV transmission rates (*γ*_*h*_) [8].

Transmission rates for low risk susceptibles engaging with high risk infected who are unaware is estimated using chronic to late stage HIV transmission rates (*γ*_*oa*_) [8]. The transmission rate is reduced by 1 *-β* for interaction with diagnosed high risk infected individuals [1]. The transmission rate for low risk susceptibles engaging with low risk infecteds, *γ*_*o*_, is assumed to be the chronic HIV transmission rates [8].

Using data from [3], we calculate that approximately 50% of individuals infected between 2006 and 2008 are diagnosed within one year [5]. In our model, newly infected susceptibles are immediately moved into the diagnosed category. We set *θ* to be 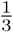 since the true number of individuals infected from 2006 to 2008 may not be known until as late as 2018. The diagnosis rate for each year thereafter is estimated as 10%, and is represented as *δ* in the model.

The growth rate of the susceptible population is taken to be 2%, comparable to the growth rate of the general U.S. population (*r*(*S*_*H*_ +*S*_*L*_)). The mortality of the susceptible population is *E*_0_, the mortality of the general U.S. population. The mortality of the infected population *E*_1_ is obtained from [5].

### 4.1. Discussion

Simulation results are shown in Fig. 2. The rate of increase for high risk diagnosed is 10%, while the rate of increase for high risk undiagnosed infected is 50%. Low risk infected increases by 12% in the 5th year. High risk undiagnosed individuals increase the fastest and remain to be the highest concern in controlling the disease.

## 5. Urban population dynamics

We consider the urban situation separately due to the higher population density, which translates into higher density sexual networks. The transmission rates of the urban population appear to be much higher than that of the general population. This effect is most likely due to increased ability for larger partner numbers than in rural or small areas. We consider the growth rate of the susceptible population to be equivalent to the national case and reduce the movement of diagnosed active individuals into the inactive pool. We modify the parameters to reflect a larger high risk susceptible population as well as a larger movement from low to high risk of the susceptibles.

Due to the larger infected population and higher diagnosis rates, we consider transmission to be more likely in high population density areas. The data on the HIV infected population in high population density areas indicates a high diagnosis rate, so we increased the number of individuals diagnosed in their first year of infection as well as a higher frequency of diagnosis in the subsequent years. The death rate in Detroit due to HIV infection is approximately 6 times larger than national [5, 13]. We incorporate this into the model by increasing the death rate of infected individuals.

### 5.1. Discussion

In the national case, active undiagnosed infected populations grow up to 50% in 5 years (Fig. 2). In the urban case, the undiagnosed active infected population quadrupled in the same time frame (Fig. 3). The diagnosed active doubled and the inactive population grows at about 25%. Overall, the undiagnosed high risk population is the main group driving the dynamics of HIV infection. For the susceptible, our model suggests that nationally the high risk susceptible population grows, but the high risk susceptible population in high population density areas decreases over time. The high risk susceptible population is clearly a concern.

**Figure 3.**
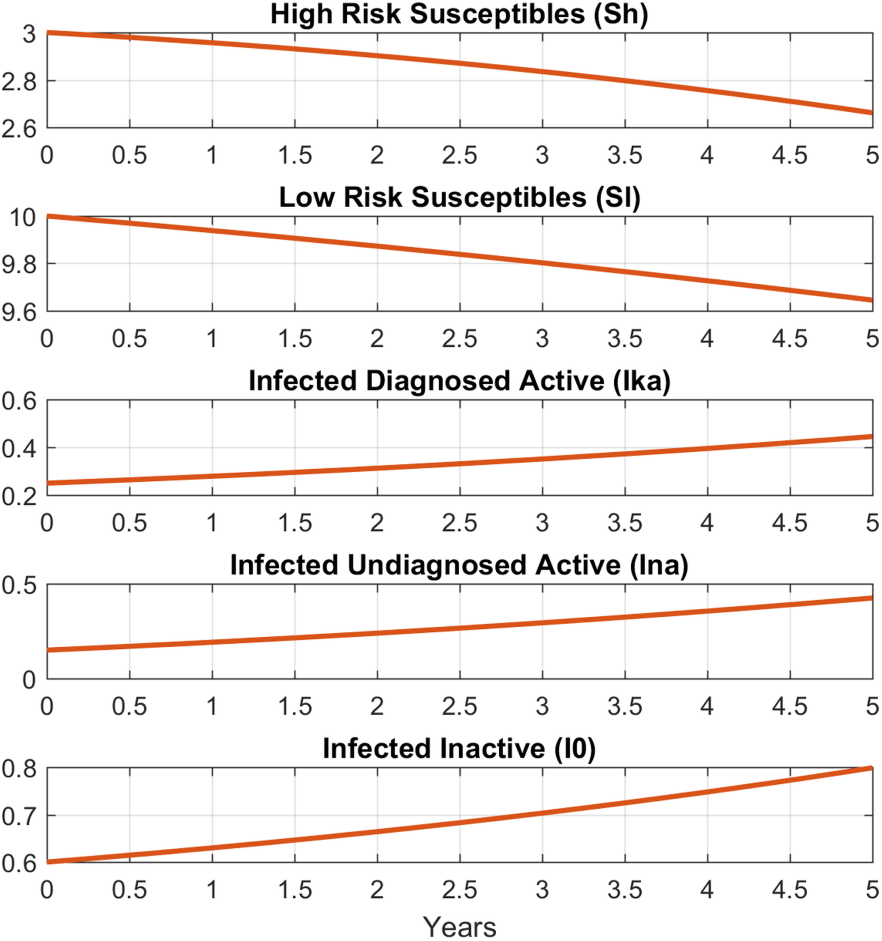
Model simulation of the urban HIV population dynamics. Parameters provided in Table 1.

**Figure 4.**
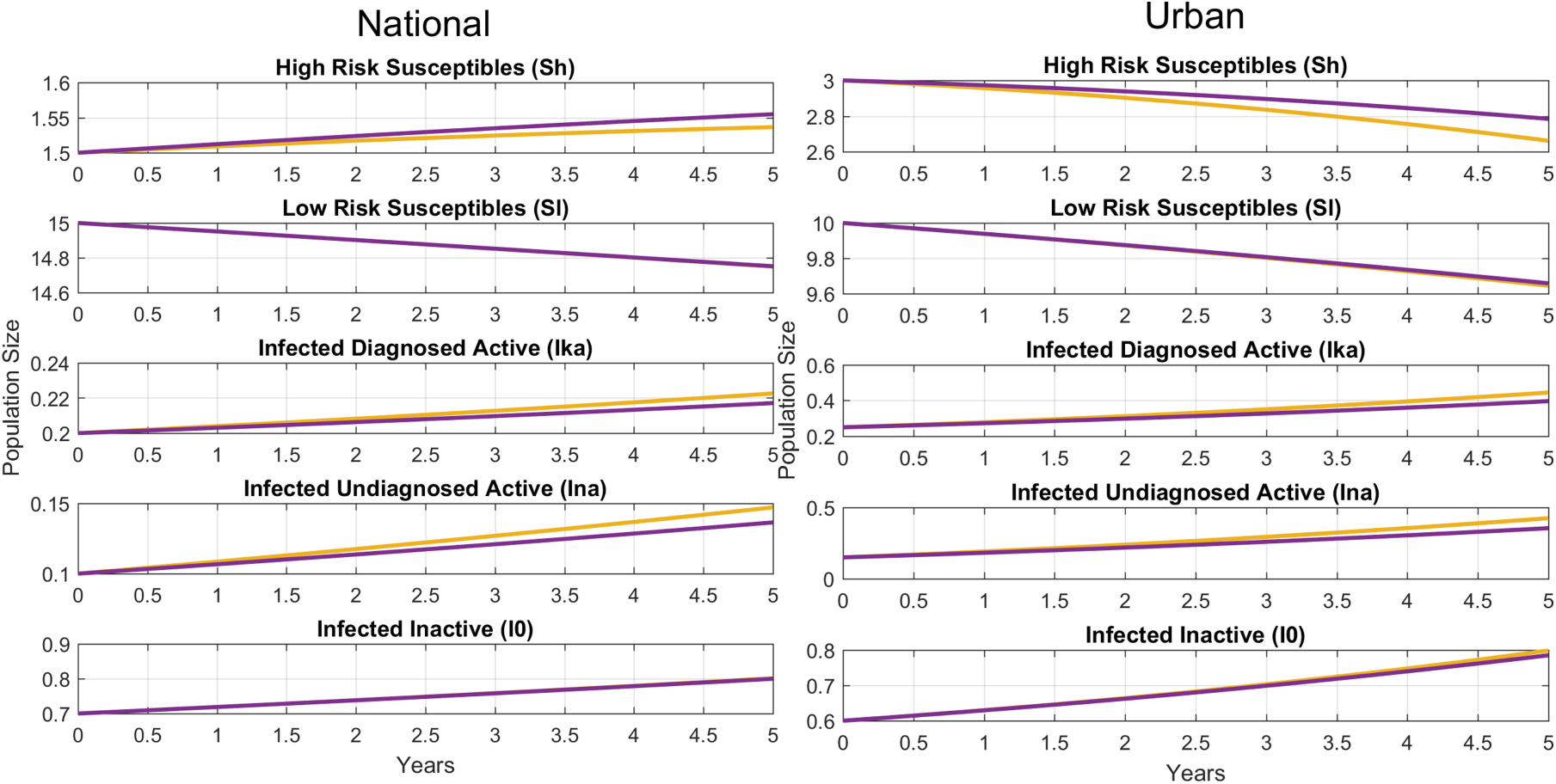
No preventative medication (yellow) compared with 20% high risk susceptibles on medication per year (purple) on the national (left) and urban (right) situations. Preventative would appear to be more effective for the population in general than in high density infective populations.

## 6. Preventative medication protocol

In this section we modify the system of equations to study the effect of pre-exposure prophylaxis (PreP). We ignore the effect of adherence as preliminary results indicate that even poor adherence (60-70%) provides protection against contraction of the disease. The removal of individuals from the pool of susceptibles due to preventative medication usage is represented by *π*. The rate of PreP usage for high risk susceptibles is *π*_*h*_ and *π*_*l*_ for low risk susceptibles. We consider the effects of different magnitudes of preventative effort, from 5% to 30% of the susceptible population yearly.

**Table 1.**
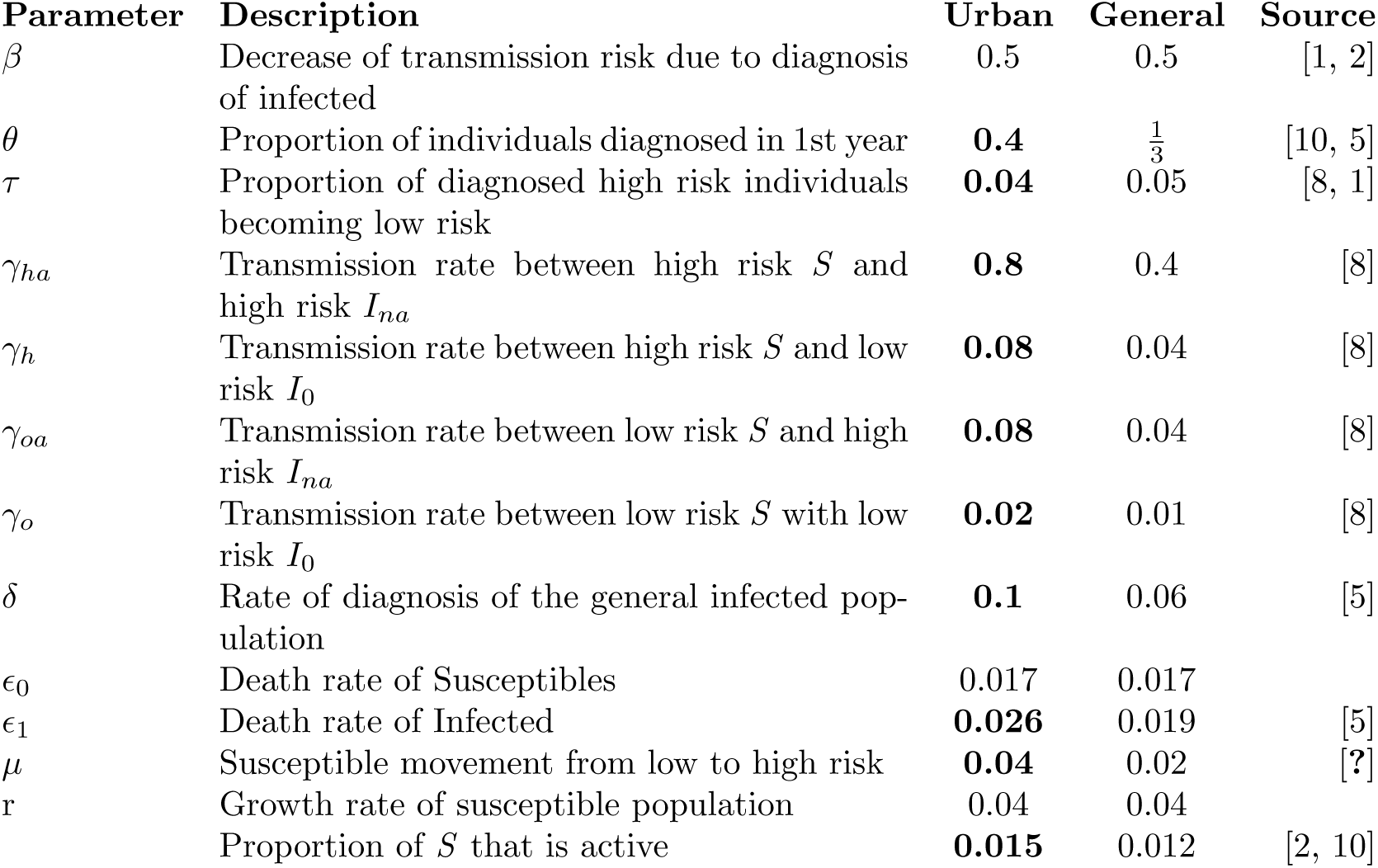
Comparison of the parameters for the Urban and General population in the US. Bolding indicated altered values. Values are taken from the literature when available, or estimated using literature relationships and proportions.

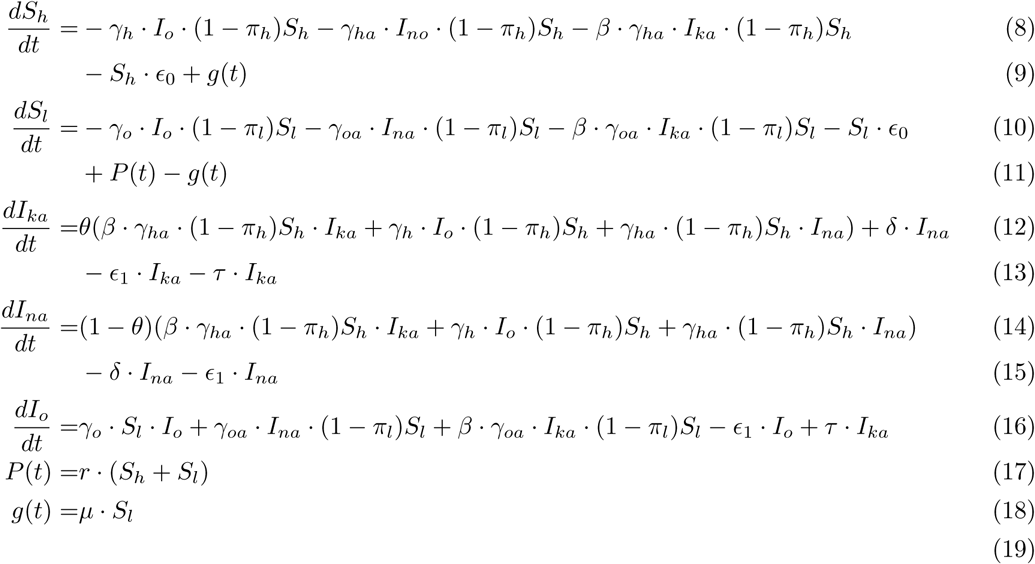

This is a control system where the *π*’s are control parameters. Ideally, one wants to vary *π* to achieve the best reduction of HIV transmission. In this paper, we take the control parameters to be constants. We wish to address time variant control parameter in the future.

### 6.1. Discussion

We find a linearly increasing reduction on the size of the active infected populations with PreP usage with both high and low risk susceptibles (Supplemental Figure 1 and 2). Our analysis suggests that targeting high risk susceptibles is far more effective than targeting low risk susceptibles or a some combination. We believe this is due to the much higher size of the low risk population, as well as reduced risk in transmission. Minimal reductions are observed in the infected inactive population. This result is to be expected since this population is considered stable.

## 7. Conclusion

When comparing the effect of *π*_*h*_ and *π*_*l*_ separately, as well as their combinations, we can see the populations are much less sensitive to changes in *π*_*l*_ (Supplemental Figure S.1 and S.2). The model suggests that targeting the low risk susceptible population would not be very effective, both in the size of transmission reduction and cost. The low risk susceptible population is estimated to be 10 times larger than the high risk population [2]. The same effect can be obtained by providing PreP to high risk susceptibles while reducing the cost by an order of magnitude. The model also suggests that minimal effect would be observed on the inactive infected population. This subpopulation includes both diagnosed and undiagnosed individuals who very rarely interact with susceptibles. We don’t expect large changes to occur in this population since they have minimal effect on the HIV epidemic.

Our results suggest that programmes targeting PreP to high risk HIV-negative individuals would have similar levels of effectiveness at the national and urban level. The model predicts 40 to 50% reductions in the size of the diagnosed and undiagnosed active infected populations after 5 years with a medication rate of 35% (Supplemental Table S.1). Previously a 44% reduction in the transmission rate was found in the original study with HIV seronegative men [14]. The increased effectiveness shown in our model is due to herd immunity. Not only are individuals taking PreP protected from HIV transmission, but they confer a degree of immunity to others within the pool of HIV-negative individuals who are at-risk.

Our study suggests that targeting PreP to high risk HIV-negative individuals provides effective protection for the general HIV-negative population, including those who are lower risk. We also find that targeting rural, smaller sexual networks can effectively curtail the growth of the HIV epidemic. Research suggests that rural individuals have higher risk profiles, as well as more difficulty accessing care [15]. The CDC has reported multiple HIV outbreaks in rural areas involving socio-economically disadvantaged individuals [16, 17], and the New York Times recently reported on a rural community with a high density of individuals dying due to lacking access to medical attention for their HIV infections [18]. We emphasize that our model makes some assumptions about the sexual network densities of high risk individuals. We expect that, in the real scenario, the benefits provided by herd immunity will increase with increasing PreP usage over the 35% we consider here.

## Acknowledgments

Thomas Le worked at the initial stage of this project. We would like to thank him for his contributions. We are grateful to NIH for providing the funding for this project.

## 9. Supplemental Information

**Figure S.1.**
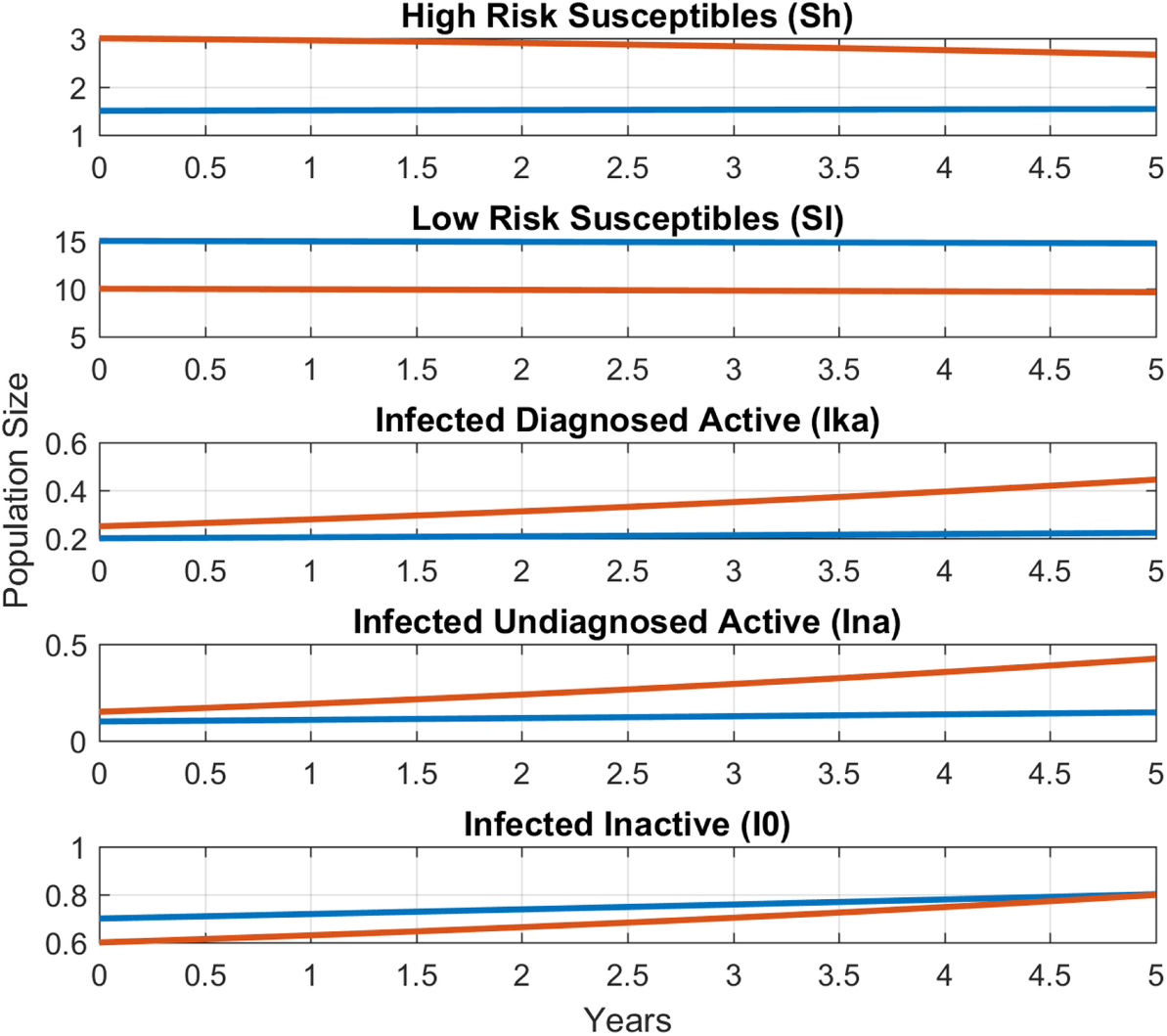
Model simulation of the urban HIV population dynamics (red) is compared to the simulation of national dynamics (blue). Parameters provided in Table 1.

**Figure S.2.**
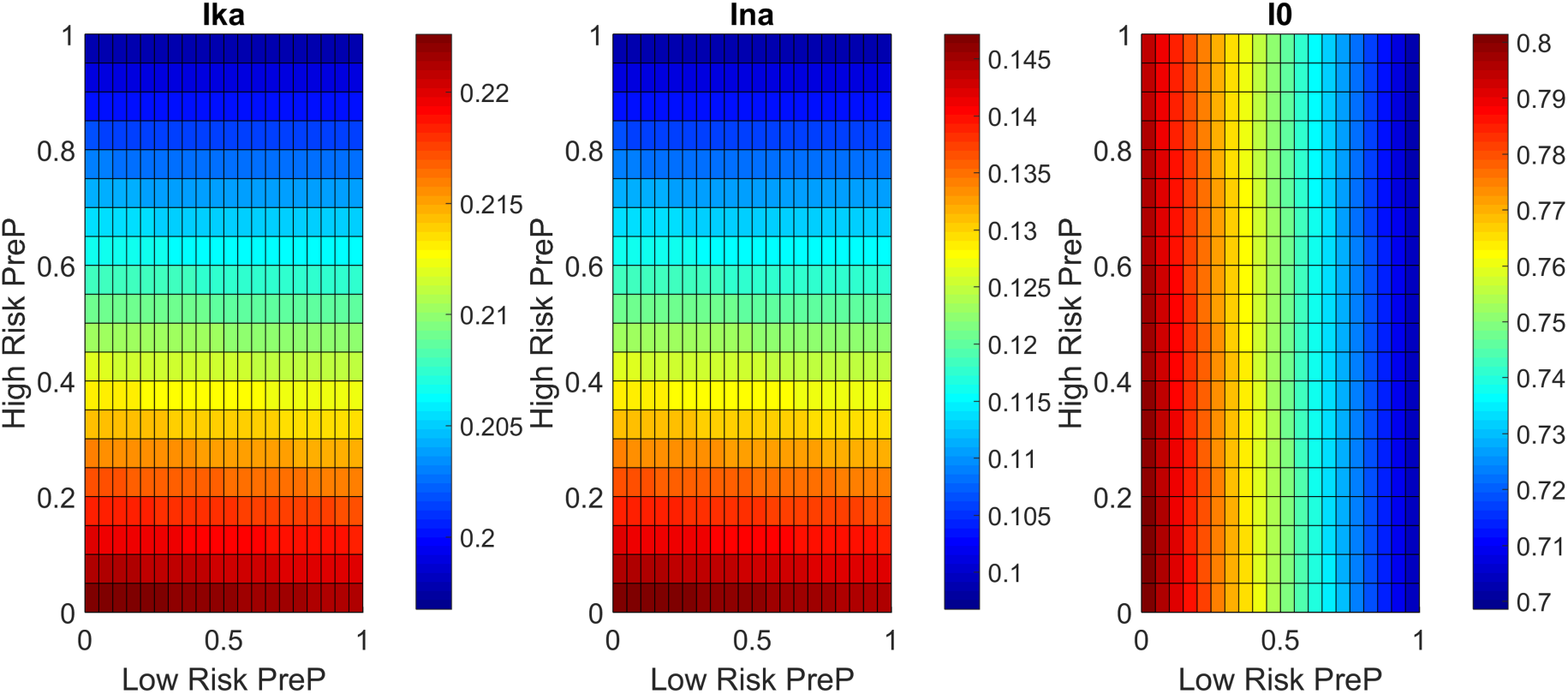
Effect of *π*_*h*_ and *π*_*l*_ on the national population dynamics after 5 years. Vertical and horizontal axes correspond to the percent coverage of PreP of the labeled group.

**Table S.1.**
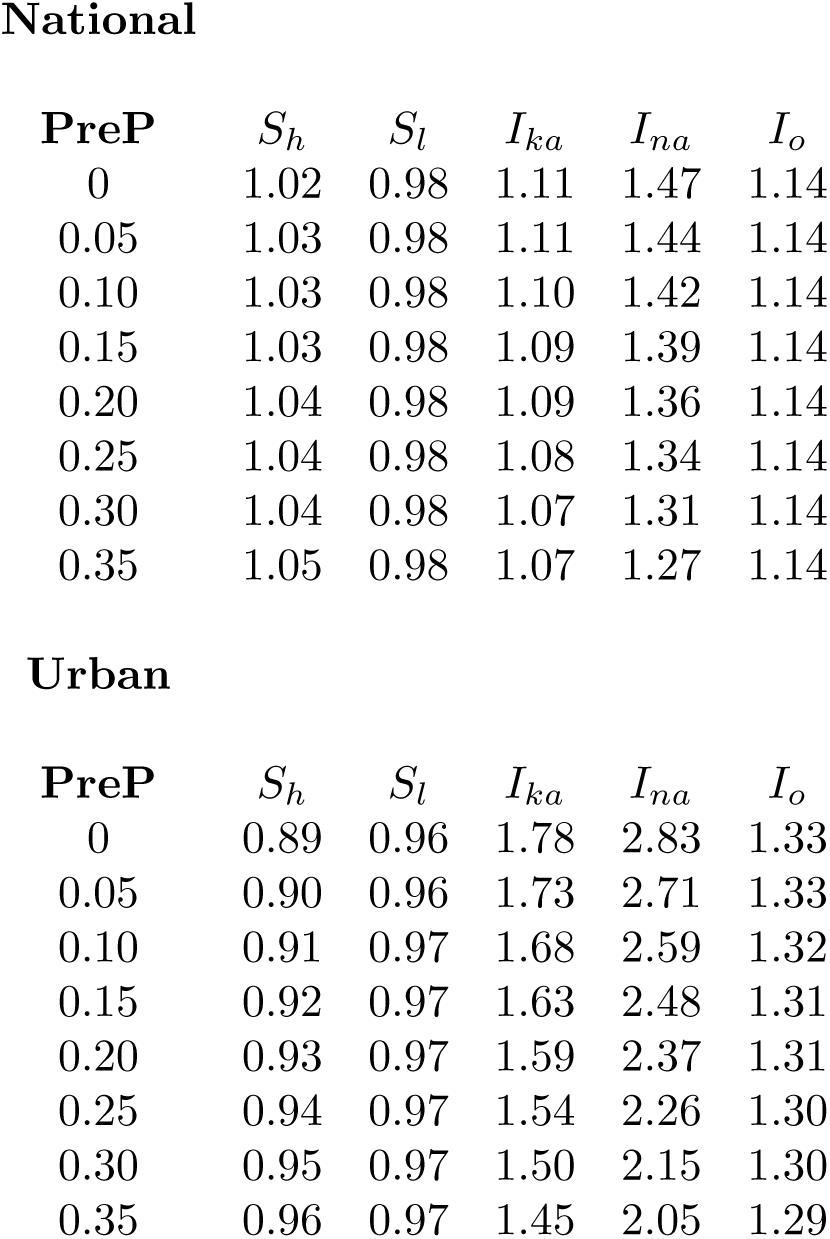
Comparison of the effect of a high risk susceptible PreP protocol on the susceptible and infected population after 5 years. A value *<* 1 indicates a reduction over 5 years, while a value *>* 1 indicates an increase.

**National**
| **PreP** | $S_h$ | $S_l$ | $I_{ka}$ | $I_{na}$ | $I_o$ |
| --- | --- | --- | --- | --- | --- |
| 0 | 1.02 | 0.98 | 1.11 | 1.47 | 1.14 |
| 0.05 | 1.03 | 0.98 | 1.11 | 1.44 | 1.14 |
| 0.10 | 1.03 | 0.98 | 1.10 | 1.42 | 1.14 |
| 0.15 | 1.03 | 0.98 | 1.09 | 1.39 | 1.14 |
| 0.20 | 1.04 | 0.98 | 1.09 | 1.36 | 1.14 |
| 0.25 | 1.04 | 0.98 | 1.08 | 1.34 | 1.14 |
| 0.30 | 1.04 | 0.98 | 1.07 | 1.31 | 1.14 |
| 0.35 | 1.05 | 0.98 | 1.07 | 1.27 | 1.14 |

| **PreP** | $S_h$ | $S_l$ | $I_{ka}$ | $I_{na}$ | $I_o$ |
| --- | --- | --- | --- | --- | --- |
| 0 | 0.89 | 0.96 | 1.78 | 2.83 | 1.33 |
| 0.05 | 0.90 | 0.96 | 1.73 | 2.71 | 1.33 |
| 0.10 | 0.91 | 0.97 | 1.68 | 2.59 | 1.32 |
| 0.15 | 0.92 | 0.97 | 1.63 | 2.48 | 1.31 |
| 0.20 | 0.93 | 0.97 | 1.59 | 2.37 | 1.31 |
| 0.25 | 0.94 | 0.97 | 1.54 | 2.26 | 1.30 |
| 0.30 | 0.95 | 0.97 | 1.50 | 2.15 | 1.30 |
| 0.35 | 0.96 | 0.97 | 1.45 | 2.05 | 1.29 |

**Figure S3.**
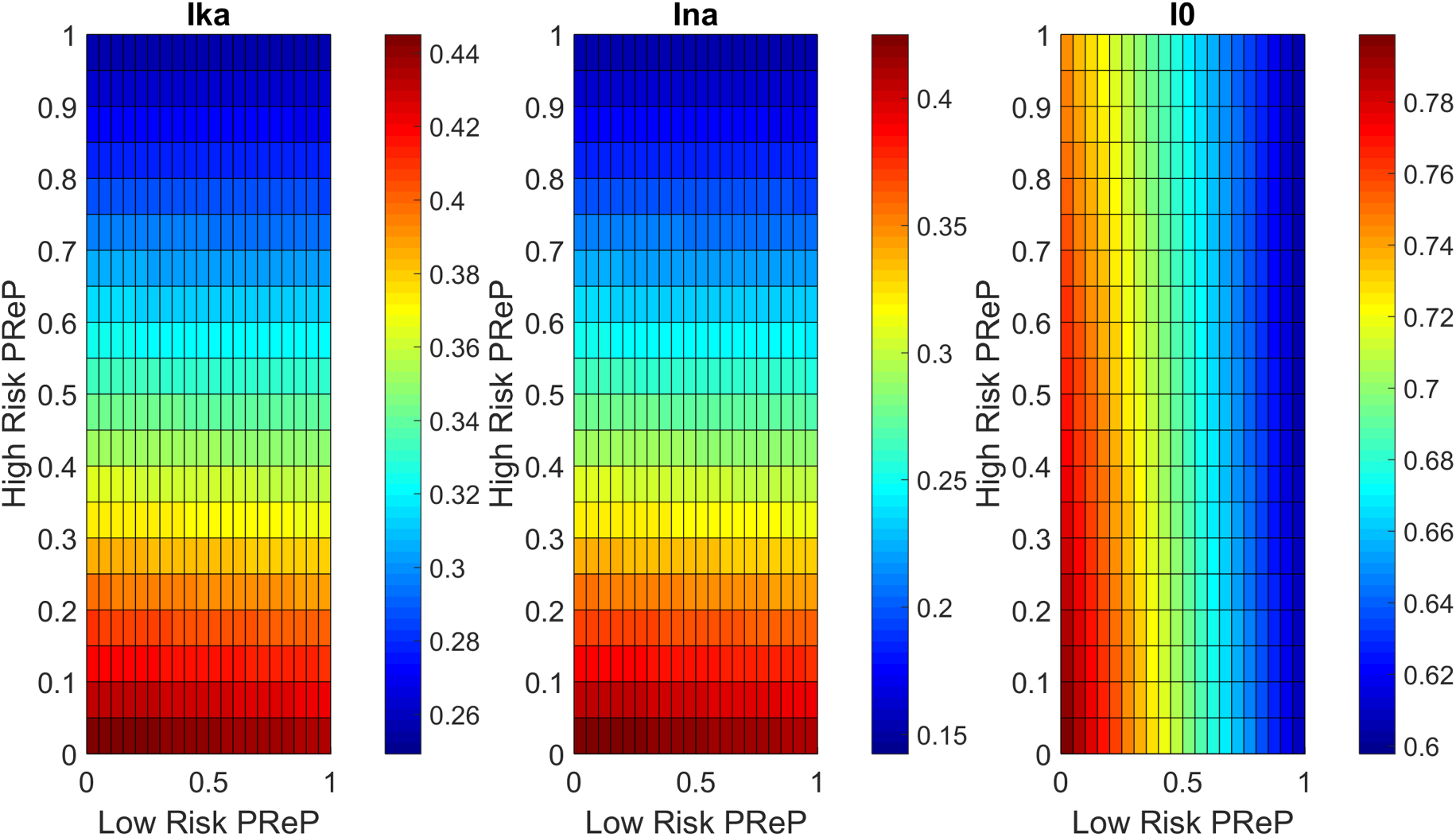
Effect of *π*_*h*_ and *π*_*l*_ on the urban population dynamics after 5 years. Vertical and horizontal axes correspond to the percent coverage of PreP of the labeled group.

